# BioNERO: an all-in-one R/Bioconductor package for comprehensive and easy biological network reconstruction

**DOI:** 10.1101/2021.04.10.439287

**Authors:** Fabricio Almeida-Silva, Thiago M. Venancio

## Abstract

**Summary:** Currently, standard network analysis workflows rely on many different packages, often requiring users to have a solid statistics and programming background. Here, we present *BioNERO,* an R package that aims to integrate all aspects of network analysis workflows, including expression data preprocessing, gene coexpression and regulatory network inference, functional analyses, and intra and interspecies network comparisons. The state-of-the-art methods implemented in *BioNERO* ensure that users can perform all analyses with a single package in a simple pipeline, without needing to learn a myriad of package-specific syntaxes. *BioNERO* offers a user-friendly framework that can be easily incorporated in systems biology pipelines.

**Availability and implementation:** The package is available at Bioconductor (http://bioconductor.org/packages/BioNERO).

## 1 Introduction

To date, several packages have been developed to infer gene coexpression networks (GCNs) and gene regulatory networks (GRN) from expression data, such as WGCNA (Langfelder and Horvath, 2008), CEMiTool (Russo *et al.*, 2018), petal (Petereit *et al.*, 2016), and minet (Meyer *et al.*, 2008). However, none of them can handle all aspects of network analysis workflows, and users are required to use other packages to build a standard analysis pipeline. Further, network inference requires a solid linear algebra and statistics background, resulting in a struggle for inexperienced researchers to properly preprocess their expression data and extract biologically meaningful information from the inferred networks.

Here, we present *BioNERO* (Biological Network Reconstruction Omnibus), an R/Bioconductor package that integrates all steps of network inference workflows in a single package. *BioNERO* uses state-of-the-art methods to preprocess expression data, infer GCNs and GRNs from expression data, analyze networks for biological interpretations, and compare networks within and across species. Additionally, *BioNERO* can be used to explore topological properties of protein-protein interaction networks, such as hub identification and community detection.

## 2 Implementation

*BioNERO* is an R package that integrates existing functionalities and introduces new ones. The input data can be common Bioconductor classes, such as SummarizedExperiment objects (Morgan *et al.*, 2020) for expression data, or basic R object classes, ensuring interoperability with other packages. Long-running functions, such as that used for Fisher’s exact tests in overrepresentation analyses, have been parallelized with BiocParallel (Morgan *et al.*, 2021) to increase speed.

### 2.1 Data preprocessing

Networks inferred from unfiltered data often do not satisfy the scale-free topology (SFT) assumption. Although this can be a property of the input data (particularly for heterogenous data sets), this issue mainly results from a lack of systematic preprocessing. In *BioNERO*, expression data are preprocessed prior to network inference to i. remove missing data; ii. remove genes with low expression across samples; iii. select genes with the highest variances (optional) and; iv. remove confounders that could introduce false-positive correlations. Count data can also be variance stabilizing transformed with DESeq2’s algorithm (Love *et al.*, 2014) to make the expression matrix approximately homoscedastic. The resulting quantile normalized expression data are adjusted for confounders based on a previously developed principal component-based method (Parsana *et al.*, 2019).

### 2.2 Gene coexpression network inference

We implemented the popular Weighted Gene Coexpression Network Analysis (WGCNA) (Langfelder and Horvath, 2008) algorithm in *BioNERO* to infer weighted networks from expression data. Users can infer three types of GCNs (signed, signed hybrid or unsigned), and pairwise gene-gene correlations can be calculated with Pearson’s r, Spearman’s ρ, or biweight midcorrelation (median-based, which is less sensible to outliers). Downstream GCN analyses in *BioNERO* include module stability evaluation, hub gene identification, functional enrichment analyses, subgraph extraction and network visualization. For all subgraph extractions, users can verify if the graphs fit the SFT, which is characteristic of real-world biological networks (Barabási *et al.*, 2011). Additionally, *BioNERO* can be used to calculate main network statistics, namely connectivity, scaled connectivity, clustering coefficient, maximum adjacency ratio, density, centralization, heterogeneity, number of cliques, diameter, betweenness, and closeness.

### 2.3 Gene regulatory network inference

Different GRN inference algorithms can be the best performers depending on the benchmark expression data set, as demonstrated by Marbach *et al.* (2012). This observation inspired the “wisdom of the crowds” principle for GRN inference, which consists in calculating average ranks for all edges across different algorithms to obtain consensus, high-confidence edges (Marbach *et al.*, 2012). Here, we implemented three widely used GRN inference algorithms: GENIE3 (Huynh-Thu *et al.*, 2010), ARACNE (Margolin *et al.*, 2006), and CLR (Faith *et al.*, 2007). However, choosing the most appropriate number of top edges to keep is a persisting bottleneck, and users often pick an arbitrary number. We implemented a method to simulate different networks by splitting the graph in *n* subgraphs, each containing the top *n*^th^ quantiles. Then, we calculate SFT fit statistics for each subgraph and select the top number of edges that leads to the best SFT fit.

### 2.4 Network comparison

GCNs inferred from different expression sets have similarities and divergences. We implemented two network comparison features in *BioNERO,* namely consensus module identification and module preservation. Consensus modules are gene modules that co-occur in networks inferred from independent expression sets, and they can be used to explore core components of the studied phenotype that are not affected by experimental effects or natural biological variation. While consensus modules identification focuses on the similarities between networks, module preservation focuses on the differences, and it can be used to explore patterns of transcriptional divergence within and across species. For interspecies comparisons, *BioNERO* can interoperate with OrthoFinder (Emms and Kelly, 2015) to analyze expression profiles at the orthogroup level.

## 3 Benchmark

A benchmark using maize (*Zea mays*) and rice (*Oryza sativa*) gene expression data obtained from Shin *et al.* (2020) is available as Supplementary Text online.

## 4 Conclusions

*BioNERO* is a novel R package that integrates all steps of network analysis pipelines, providing users with a simple framework for GCN and GRN inference from expression data. This package can be easily integrated in systems biology pipelines and will likely accelerate biological network analysis projects.

## Acknowledgements

This work was supported by Fundação Carlos Chagas Filho de Amparo à Pesquisa do Estado do Rio de Janeiro (FAPERJ; grants E-26/203.309/2016 and E-26/203.014/2018), Coordenação de Aperfeiçoamento de Pessoal de Nível Superior - Brasil (CAPES; Finance Code 001), and Conselho Nacional de Desenvolvimento Científico e Tecnológico. The funding agencies had no role in the design of the study and collection, analysis, and interpretation of data and in writing.

Conflicts of interest: none declared.

## REFERENCES

Barabási,A.-L. et al. (2011) Hierarchical Organization of Modularity in Complex Networks. Science (80-.)., 297, 46–65.

Emms,D.M. and Kelly,S. (2015) OrthoFinder: solving fundamental biases in whole genome comparisons dramatically improves orthogroup inference accuracy. Genome Biol., 16, 1–14.

Faith,J.J. et al. (2007) Large-scale mapping and validation of Escherichia coli transcriptional regulation from a compendium of expression profiles. PLoS Biol., 5, 0054–0066.

Huynh-Thu,V.A. et al. (2010) Inferring regulatory networks from expression data using tree-based methods. PLoS One, 5, 1–10.

Langfelder,P. and Horvath,S. (2008) WGCNA: an R package for weighted correlation network analysis. BMC Bioinformatics, 9, 559.

Love,M.I. et al. (2014) Moderated estimation of fold change and dispersion for RNA-seq data with DESeq2. Genome Biol., 15, 1–21.

Marbach,D. et al. (2012) Wisdom of crowds for robust gene network inference. Nat. Methods, 9, 796–804.

Margolin,A.A. et al. (2006) ARACNE: An algorithm for the reconstruction of gene regulatory networks in a mammalian cellular context. BMC Bioinformatics, 7, 1–15.

Meyer,P.E. et al. (2008) Minet: A r/bioconductor package for inferring large transcriptional networks using mutual information. BMC Bioinformatics, 9, 1–10.

Morgan,M. et al. (2021) BiocParallel: Bioconductor facilities for parallel evaluation.

Morgan,M. et al. (2020) SummarizedExperiment: SummarizedExperiment container.

Parsana,P. et al. (2019) Addressing confounding artifacts in reconstruction of gene co-expression networks. Genome Biol., 20, 94.

Petereit,J. et al. (2016) petal: Co-expression network modelling in R. BMC Syst. Biol., 10, 51.

Russo,P.S.T. et al. (2018) CEMiTool: a Bioconductor package for performing comprehensive modular co-expression analyses. BMC Bioinformatics, 19, 56.

Shin,J. et al. (2020) A network-based comparative framework to study conservation and divergence of proteomes in plant phylogenies. Nucleic Acids Res., 1–23.

