## Supplementary Text for "BioNERO: an all-in-one R/Bioconductor package for comprehensive and easy biological network reconstruction"

[1]Universidade Estadual do Norte Fluminense Darcy Ribeiro

**April 2021**

### 1 Data description

The benchmark expression data correspond to maize and rice gene expression data normalized in TPM, which were obtained from Shin et al. (2020). Maize transcription factors were downloaded from PlantTFDB 4.0 (Jin et al. 2017). Functional gene annotation and orthogroups between maize and rice were downloaded from PLAZA 4.0 Monocots (Van Bel et al. 2018). The objects we have pre-loaded are:

- **maize_se:** SummarizedExperiment object with maize gene expression data.
- **rice_se:** SummarizedExperiment object with rice gene expression data.
- **zma_annotation:** Data frame with Interpro and Gene Ontology annotation for maize genes.
- **zma_tfs:** Data frame with gene IDs of transcription factors and their families.
- **orthogroups:** Data frame with orthogroups between maize and rice.

```
library(BioNERO)
##
set.seed(1) # for reproducibility
```

### 2 Data preprocessing

Here, we will remove genes with median expression levels <5.

```
maize_exp <- exp_preprocess(maize_se, min_exp = 5)
## Number of removed samples: 4
```

### 3 Exploratory analysis

To start with, let's explore sample clustering with a heatmap of hierarchically clustered samples. Then, we will perform a principal component analysis (PCA) to observe clustering patterns.

```
# Heatmap of sample correlations
plot_heatmap(maize_exp, type = "samplecor", show_colnames=FALSE)
```

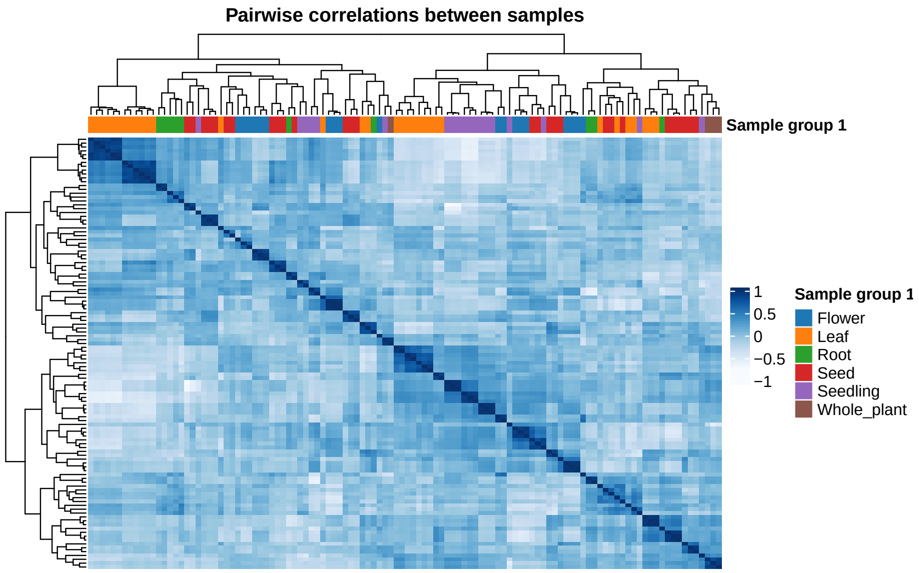

```
# PCA plot
plot_PCA(maize_exp, size=3)
```

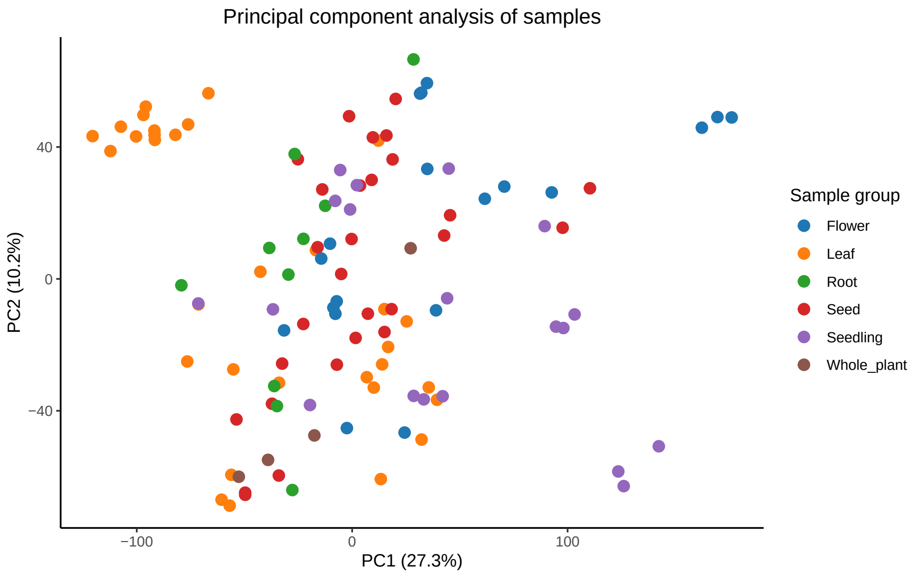

As the data set is highly heterogeneous, with samples from different developmental stages and stress conditions for the same tissues, we did not expect a perfect clustering by tissue. Hence, the patterns in the plots match what we expected.

### 4 GCN inference and analysis

First, we need to calculate the optimal $\beta$ power based on the scale-free topology fit. Then, we use this power to infer the GCN.

```
# Calculate SFT fit for beta powers
sft <- SFT_fit(maize_exp, cor_method = "pearson")
##    Power SFT.R.sq  slope truncated.R.sq mean.k. median.k. max.k.
## 1      3   0.4070  1.660          0.908  3470.0    3520.0   5600
## 2      4   0.0658  0.447          0.773  2330.0    2320.0   4370
## 3      5   0.0457 -0.312          0.702  1620.0    1560.0   3500
## 4      6   0.2720 -0.781          0.730  1150.0    1060.0   2850
## 5      7   0.4340 -1.060          0.770   841.0     738.0   2360
## 6      8   0.5360 -1.230          0.805   627.0     520.0   1980
## 7      9   0.5960 -1.350          0.829   476.0     373.0   1670
## 8     10   0.6360 -1.450          0.845   367.0     269.0   1430
## 9     11   0.6760 -1.500          0.869   287.0     197.0   1230
## 10    12   0.7060 -1.530          0.888   228.0     145.0   1060
## 11    13   0.7330 -1.550          0.904   182.0     109.0    925
## 12    14   0.7590 -1.560          0.919   147.0      82.0    809
## 13    15   0.7740 -1.580          0.928   120.0      62.3    710
## 14    16   0.7880 -1.600          0.936    98.9      47.9    626
## 15    17   0.8000 -1.600          0.942    82.0      37.3    554
## 16    18   0.8130 -1.600          0.950    68.4      29.3    492
## 17    19   0.8140 -1.620          0.953    57.4      23.2    441
## 18    20   0.8180 -1.640          0.956    48.4      18.5    396

# Visual diagnostics
sft$plot
```

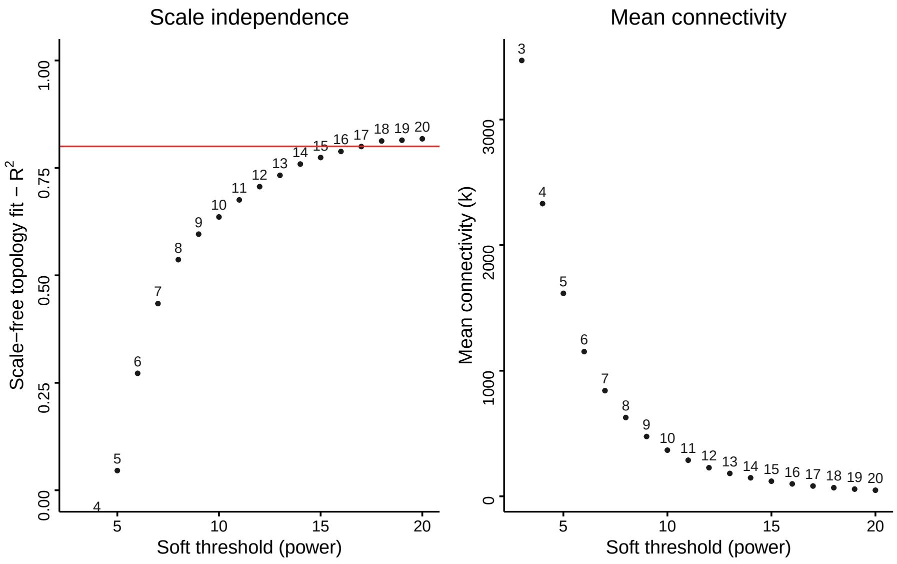

```
power <- sft$power

# Infer GCN
gcn <- exp2gcn(maize_exp, SFTpower = power, cor_method = "pearson")
## ..connectivity..
## ..matrix multiplication (system BLAS)..
## ..normalization..
```

```
## ..done.
```

Let's see the number of genes per module.

```
plot_ngenes_per_module(gcn)
```

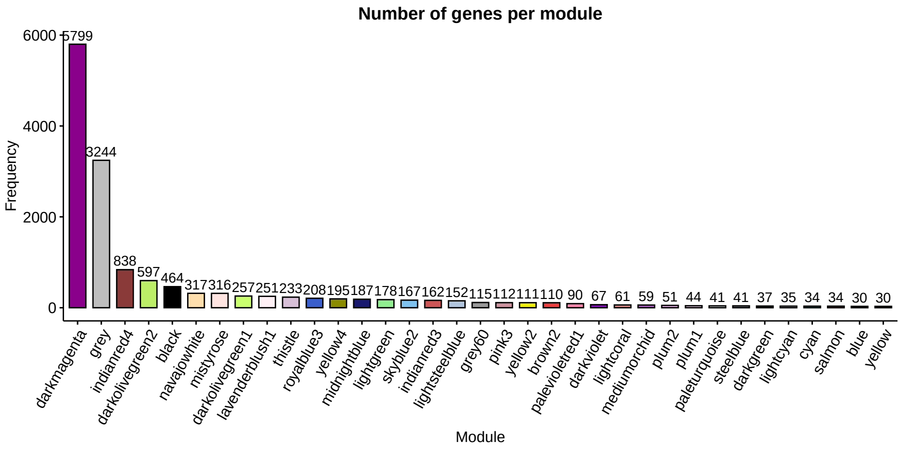

Now, we will perform an enrichment analysis to see if any of these modules is enriched in protein domains or GO terms. As background, we will use all genes in the network.

```
head(zma_annotation)
##             Gene
## 1 Zm00001d000001
## 2 Zm00001d000001
## 3 Zm00001d000001
## 4 Zm00001d000001
## 5 Zm00001d000001
## 6 Zm00001d000001
##                                                                                       GO
## 1 oxidoreductase activity, acting on diphenols and related substances as donors, oxygen as acceptor
## 2 oxidoreductase activity, acting on diphenols and related substances as donors, oxygen as acceptor
## 3 oxidoreductase activity, acting on diphenols and related substances as donors, oxygen as acceptor
## 4 oxidoreductase activity, acting on diphenols and related substances as donors, oxygen as acceptor
## 5                                                                        catalytic activity
## 6                                                                        catalytic activity
##                                    Interpro
## 1         Tyrosinase copper-binding domain
## 2           Polyphenol oxidase, C-terminal
## 3       Polyphenol oxidase, central domain
## 4 Uncharacterised domain, di-copper centre
## 5         Tyrosinase copper-binding domain
## 6           Polyphenol oxidase, C-terminal

# Perform the enrichment analysis for all modules (1 thread only)
enrich <- module_enrichment(net = gcn,
                            background_genes = rownames(maize_exp),
                            annotation = zma_annotation)
## Enrichment analysis for module black...
```

```
## Enrichment analysis for module blue...
## Enrichment analysis for module brown2...
## Enrichment analysis for module cyan...
## Enrichment analysis for module darkgreen...
## Enrichment analysis for module darkmagenta...
## Enrichment analysis for module darkolivegreen1...
## Enrichment analysis for module darkolivegreen2...
## Enrichment analysis for module darkviolet...
## Enrichment analysis for module grey60...
## Enrichment analysis for module indianred3...
## Enrichment analysis for module indianred4...
## Enrichment analysis for module lavenderblush1...
## Enrichment analysis for module lightcoral...
## Enrichment analysis for module lightcyan...
## Enrichment analysis for module lightgreen...
## Enrichment analysis for module lightsteelblue...
## Enrichment analysis for module mediumorchid...
## Enrichment analysis for module midnightblue...
## Enrichment analysis for module mistyrose...
## Enrichment analysis for module navajowhite...
## Enrichment analysis for module paleturquoise...
## Enrichment analysis for module palevioletred1...
## Enrichment analysis for module pink3...
## Enrichment analysis for module plum1...
## Enrichment analysis for module plum2...
## Enrichment analysis for module royalblue3...
## Enrichment analysis for module salmon...
## Enrichment analysis for module skyblue2...
## Enrichment analysis for module steelblue...
## Enrichment analysis for module thistle...
## Enrichment analysis for module yellow...
## Enrichment analysis for module yellow2...
## Enrichment analysis for module yellow4...
nrow(enrich)
## [1] 2865
```

As the enrichment analysis returned modules enriched in several protein domains and GO terms, we will display only the top 3 results for each module and annotation category.

```
# Show top 3 results without unadjusted P-value and gene IDs
library(tidyverse)
enrich %>%
  select(!c(GeneID, pval)) %>%
  group_by(Module, Category) %>%
  slice_min(padj, n = 3, with_ties = FALSE) %>%
  print(n=Inf)
## # A tibble: 157 x 6
## # Groups:   Module, Category [55]
##     TermID                              genes   all      padj Category Module
##     <chr>                               <dbl> <dbl>     <dbl> <chr>    <chr>
##   1 ribosomal subunit                     171   260 4.77e-198 GO       black
```

```
## 2 cytosolic ribosome                     186   337 2.23e-196 GO       black
## 3 cytosolic part                         187   351 1.04e-193 GO       black
## 4 Translation protein SH3-like domain     19    33 7.40e- 17 Interpro black
## 5 Ribosomal protein L2 domain 2           15    28 2.01e- 12 Interpro black
## 6 Zinc-binding ribosomal protein          11    16 2.19e- 10 Interpro black
## 7 cell periphery                          49  2342 9.22e-  9 GO       brown2
## 8 cell wall                               25   626 1.12e-  8 GO       brown2
## 9 external encapsulating structure        25   626 1.12e-  8 GO       brown2
## 10 Tubulin                                 6    26 1.94e-  4 Interpro brown2
## 11 Tubulin/FtsZ, GTPase domain             6    30 2.44e-  4 Interpro brown2
## 12 Tubulin/FtsZ, 2-layer sandwich dom~     5    18 3.51e-  4 Interpro brown2
## 13 photosynthetic membrane                28   384 1.70e- 35 GO       cyan
## 14 thylakoid part                         28   393 1.70e- 35 GO       cyan
## 15 chloroplast thylakoid membrane         27   351 7.64e- 35 GO       cyan
## 16 Chlorophyll A-B binding protein, p~    10    20 1.16e- 18 Interpro cyan
## 17 Chlorophyll A-B binding protein        10    22 2.02e- 18 Interpro cyan
## 18 Chlorophyll a/b binding protein do~    10    32 1.32e- 16 Interpro cyan
## 19 nucleic acid metabolic process       1050  1924 1.10e- 42 GO       darkmage~
## 20 RNA metabolic process                 887  1645 1.66e- 32 GO       darkmage~
## 21 RNA processing                        316   493 1.02e- 25 GO       darkmage~
## 22 Armadillo-like helical                175   263 2.55e- 15 Interpro darkmage~
## 23 Armadillo-type fold                   176   278 1.79e- 12 Interpro darkmage~
## 24 Zinc finger, FYVE/PHD-type             75    95 8.80e- 12 Interpro darkmage~
## 25 unfolded protein binding               12    91 4.75e-  4 GO       darkoliv~
## 26 protein folding                        14   164 4.55e-  3 GO       darkoliv~
## 27 DnaJ domain, conserved site            10    47 4.12e-  5 Interpro darkoliv~
## 28 Chaperone DnaJ, C-terminal              6    21 1.92e-  3 Interpro darkoliv~
## 29 DnaJ domain                            11    99 1.92e-  3 Interpro darkoliv~
## 30 endomembrane system                   222  1541 1.07e- 66 GO       darkoliv~
## 31 Golgi apparatus                       167   895 3.36e- 64 GO       darkoliv~
## 32 vesicle-mediated transport             70   295 5.11e- 31 GO       darkoliv~
## 33 Longin-like domain                     15    38 4.67e-  8 Interpro darkoliv~
## 34 Nucleotide-diphospho-sugar transfe~    23   106 1.00e-  7 Interpro darkoliv~
## 35 Nonaspanin (TM9SF)                      9    18 2.07e-  5 Interpro darkoliv~
## 36 intracellular distribution of mito~     2     2 2.32e-  2 GO       darkviol~
## 37 mitochondrial fusion                    2     2 2.32e-  2 GO       darkviol~
## 38 mitochondrion distribution              2     2 2.32e-  2 GO       darkviol~
## 39 nucleolus                              36   574 8.10e- 14 GO       indianre~
## 40 ncRNA metabolic process                21   249 1.06e-  9 GO       indianre~
## 41 nuclear lumen                          37   861 1.06e-  9 GO       indianre~
## 42 Chaperone tailless complex polypep~     6    14 1.42e-  5 Interpro indianre~
## 43 Chaperonin TCP-1, conserved site        6    14 1.42e-  5 Interpro indianre~
## 44 WD40/YVTN repeat-like-containing d~    16   256 4.90e-  5 Interpro indianre~
## 45 endomembrane system                   141  1541 2.06e-  5 GO       indianre~
## 46 establishment of localization         136  1469 2.06e-  5 GO       indianre~
## 47 localization                          142  1561 2.06e-  5 GO       indianre~
## 48 protein kinase activity                41   629 5.09e- 10 GO       lavender~
## 49 protein phosphorylation                41   631 5.09e- 10 GO       lavender~
## 50 phosphotransferase activity, alcoh~    41   757 1.17e-  7 GO       lavender~
## 51 Protein kinase-like domain             36   559 3.79e-  8 Interpro lavender~
## 52 Protein kinase domain                  33   513 1.63e-  7 Interpro lavender~
```

```
## 53 Protein kinase, ATP binding site       26   343 3.19e-  7 Interpro lavender~
## 54 monocarboxylic acid metabolic proc~    27   363 9.38e- 24 GO       lightcor~
## 55 fatty acid biosynthetic process        15    99 1.55e- 16 GO       lightcor~
## 56 fatty acid metabolic process           17   159 1.55e- 16 GO       lightcor~
## 57 Biotin/lipoyl attachment                6    18 3.21e-  7 Interpro lightcor~
## 58 Single hybrid motif                     6    19 3.21e-  7 Interpro lightcor~
## 59 E3-binding domain                       4     9 6.87e-  5 Interpro lightcor~
## 60 1,4-beta-D-xylan synthase activity      4    19 2.03e-  4 GO       lightcyan
## 61 glucuronoxylan biosynthetic process     4    19 2.03e-  4 GO       lightcyan
## 62 glucuronoxylan metabolic process        4    19 2.03e-  4 GO       lightcyan
## 63 Glycosyl transferase, family 43         3    10 9.06e-  3 Interpro lightcyan
## 64 chloroplast                            65  2308 1.09e- 11 GO       lightste~
## 65 plastid                                65  2413 4.94e- 11 GO       lightste~
## 66 chloroplast thylakoid membrane         16   351 1.94e-  3 GO       lightste~
## 67 mitochondrial protein complex          12   130 8.73e- 10 GO       mediumor~
## 68 inner mitochondrial membrane prote~    11   124 5.27e-  9 GO       mediumor~
## 69 mitochondrial membrane part            12   160 5.27e-  9 GO       mediumor~
## 70 Myristoyl-CoA:protein N-myristoylt~     2     3 3.65e-  2 Interpro mediumor~
## 71 Myristoyl-CoA:protein N-myristoylt~     2     3 3.65e-  2 Interpro mediumor~
## 72 Myristoyl-CoA:protein N-myristoylt~     2     3 3.65e-  2 Interpro mediumor~
## 73 chloroplast                           149  2308 4.56e- 81 GO       midnight~
## 74 plastid                               150  2413 7.28e- 80 GO       midnight~
## 75 plastid part                          124  1390 3.19e- 78 GO       midnight~
## 76 L28p-like                               4     7 4.89e-  3 Interpro midnight~
## 77 Peroxiredoxin, AhpC-type                3     3 4.89e-  3 Interpro midnight~
## 78 Rhodanese-like domain                   5    18 4.89e-  3 Interpro midnight~
## 79 chromosome organization                53   382 3.99e- 24 GO       mistyrose
## 80 DNA conformation change                28    93 3.37e- 21 GO       mistyrose
## 81 DNA metabolic process                  49   408 1.18e- 19 GO       mistyrose
## 82 Kinesin motor domain                   15    38 4.37e- 12 Interpro mistyrose
## 83 Kinesin motor domain, conserved si~    12    26 1.84e- 10 Interpro mistyrose
## 84 MCM N-terminal domain                   8    10 3.77e-  9 Interpro mistyrose
## 85 GLABROUS1 enhancer-binding protein~     5     9 3.29e-  3 Interpro navajowh~
## 86 F-box domain                           13   119 5.39e-  3 Interpro navajowh~
## 87 antifungal innate immune response       2     4 3.45e-  2 GO       paleturq~
## 88 defense response to fungus              6   244 3.45e-  2 GO       paleturq~
## 89 diphosphoinositol-pentakisphosphat~     2     5 3.45e-  2 GO       paleturq~
## 90 AP2/ERF domain                          4    35 1.59e-  2 Interpro paleturq~
## 91 DNA-binding domain                      4    45 2.21e-  2 Interpro paleturq~
## 92 cellular response to alkaline pH        4     7 1.21e-  4 GO       paleviol~
## 93 cellular response to pH                 4     7 1.21e-  4 GO       paleviol~
## 94 regulation of cellular response to~     4     7 1.21e-  4 GO       paleviol~
## 95 CO/COL/TOC1, conserved site             4    11 2.60e-  3 Interpro paleviol~
## 96 Tify domain                             4    14 3.89e-  3 Interpro paleviol~
## 97 protein kinase activity                25   629 3.06e-  8 GO       pink3
## 98 protein phosphorylation                25   631 3.06e-  8 GO       pink3
## 99 cellular protein modification proc~    32  1262 8.80e-  7 GO       pink3
## 100 Protein kinase domain                 25   513 2.41e- 10 Interpro pink3
## 101 Protein kinase-like domain            26   559 2.41e- 10 Interpro pink3
## 102 Protein kinase, ATP binding site      19   343 2.48e-  8 Interpro pink3
## 103 structural constituent of ribosome    26   398 4.33e- 26 GO       plum1
```

```
## 104 intracellular ribonucleoprotein co~    28   626 1.42e- 24 GO       plum1
## 105 ribonucleoprotein complex              28   626 1.42e- 24 GO       plum1
## 106 Ribosomal protein L7Ae/L30e/S12e/G~     4    15 5.76e-  4 Interpro plum1
## 107 50S ribosomal protein L30e-like          4    19 8.11e-  4 Interpro plum1
## 108 Ribosomal protein S14                    3     6 1.02e-  3 Interpro plum1
## 109 DNA packaging complex                    6    82 7.39e-  4 GO       plum2
## 110 nucleosome                               6    81 7.39e-  4 GO       plum2
## 111 protein-DNA complex assembly             6    79 7.39e-  4 GO       plum2
## 112 Histone H5                               4     5 3.97e-  6 Interpro plum2
## 113 Linker histone H1/H5, domain H15         5    16 5.41e-  6 Interpro plum2
## 114 AT hook, DNA-binding motif               5    41 5.80e-  4 Interpro plum2
## 115 plastid                                124  2413 1.03e- 41 GO       royalblu~
## 116 chloroplast                            121  2308 1.90e- 41 GO       royalblu~
## 117 plastid part                            84  1390 1.35e- 29 GO       royalblu~
## 118 Mitochondrial inner membrane trans~     5    18 8.25e-  3 Interpro royalblu~
## 119 Trigger factor                           3     3 8.25e-  3 Interpro royalblu~
## 120 Trigger factor, ribosome-binding, ~     3     3 8.25e-  3 Interpro royalblu~
## 121 cellular response to phosphate sta~     9   114 1.87e-  8 GO       salmon
## 122 phosphate ion homeostasis                6    22 1.87e-  8 GO       salmon
## 123 trivalent inorganic anion homeosta~     6    22 1.87e-  8 GO       salmon
## 124 SPX domain-containing protein            4     4 1.48e-  7 Interpro salmon
## 125 SPX domain                               4     7 2.57e-  6 Interpro salmon
## 126 Calcineurin-like phosphoesterase d~     6    50 2.95e-  6 Interpro salmon
## 127 chloroplast thylakoid membrane          18   351 2.23e-  4 GO       skyblue2
## 128 photosynthetic membrane                 19   384 2.23e-  4 GO       skyblue2
## 129 thylakoid membrane                      19   382 2.23e-  4 GO       skyblue2
## 130 UbiB domain                              5    17 6.13e-  3 Interpro skyblue2
## 131 DNA packaging complex                   27    82 1.64e- 49 GO       steelblue
## 132 nucleosome                              27    81 1.64e- 49 GO       steelblue
## 133 protein-DNA complex                     27    90 2.17e- 48 GO       steelblue
## 134 Histone-fold                            27    86 1.18e- 48 Interpro steelblue
## 135 Histone H2A/H2B/H3                      22    48 4.98e- 43 Interpro steelblue
## 136 Histone H2B                             10    14 1.80e- 20 Interpro steelblue
## 137 chloroplast                            187  2308 6.44e-104 GO       thistle
## 138 plastid                                187  2413 1.26e-100 GO       thistle
## 139 chloroplast part                       149  1365 1.09e- 92 GO       thistle
## 140 Aminoacyl-tRNA synthetase, class II      9    26 8.37e-  7 Interpro thistle
## 141 Peptidase S1C                            4     8 1.27e-  2 Interpro thistle
## 142 Aminoacyl-tRNA synthetase, class I~     4    10 2.40e-  2 Interpro thistle
## 143 Aspartate/other aminotransferase         2     3 2.48e-  2 Interpro yellow
## 144 Pyridoxal phosphate-dependent tran~     4    65 2.48e-  2 Interpro yellow
## 145 Pyridoxal phosphate-dependent tran~     4    61 2.48e-  2 Interpro yellow
## 146 RNA processing                          17   493 1.32e-  3 GO       yellow2
## 147 RNA metabolic process                   31  1645 3.95e-  3 GO       yellow2
## 148 intracellular organelle lumen           21   918 4.61e-  3 GO       yellow2
## 149 Pentatricopeptide repeat                19    82 8.22e- 20 Interpro yellow2
## 150 Tetratricopeptide-like helical dom~    16   252 2.10e-  7 Interpro yellow2
## 151 Carbamoyl-phosphate synthase large~     2     2 2.05e-  2 Interpro yellow2
## 152 chloroplast part                        69  1365 3.41e- 20 GO       yellow4
## 153 plastid part                            70  1390 3.41e- 20 GO       yellow4
## 154 chloroplast                             86  2308 7.74e- 18 GO       yellow4
```

```
## 155 Protein of unknown function DUF2358     4     6 1.63e-  3 Interpro yellow4
## 156 Thioredoxin-like fold                  13   173 1.63e-  3 Interpro yellow4
## 157 FAD/NAD(P)-binding domain               8    69 6.05e-  3 Interpro yellow4
```

Since we now have some information on which biological processes are associated to each module, we can correlate module eigengenes (MEs) to tissues. MEs are each module's first principal component, which summarize their expression levels. Thus, if a module eigengene is positively correlated with a particular tissue, we can assume that this module contains genes whose expression levels increase in this particular tissue.

```
me_tissue_cor <- module_trait_cor(exp = maize_exp,
                                  MEs = gcn$MEs,
                                  cor_method = "pearson")
```

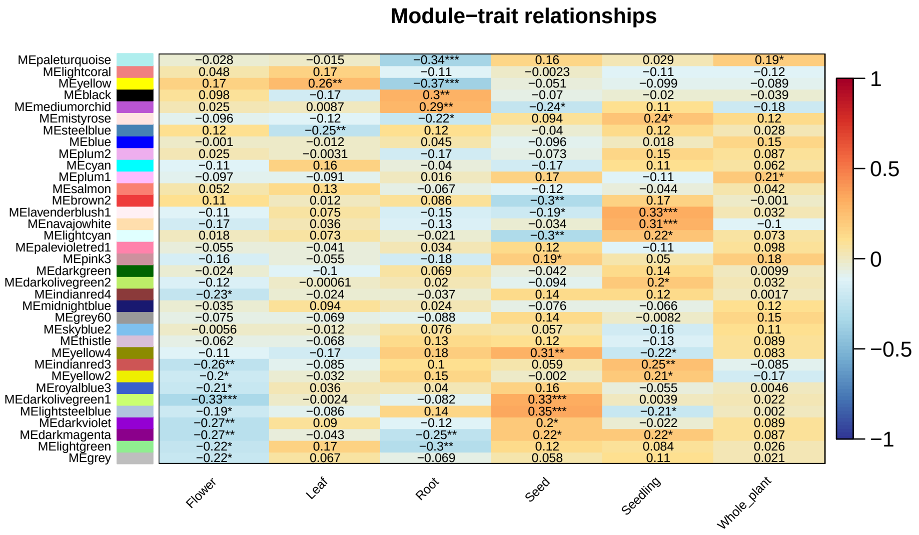

We can see that the module *black* has a significant positive correlation with root samples. Let's visualize it with a line plot.

```
plot_expression_profile(exp = maize_exp, net = gcn,
                        plot_module=TRUE, modulename="black")
```

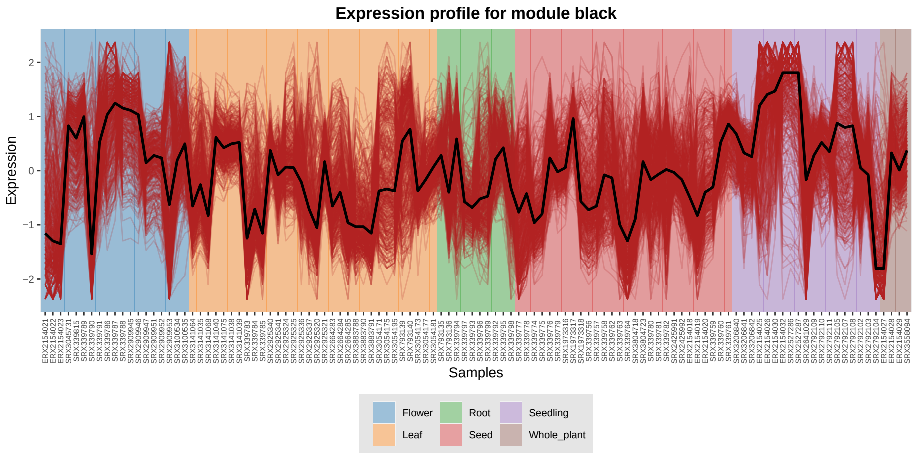

After identifying modules and their associated biological processes, we can pinpoint central genes in each module that have the highest number of connections, the so-called **intramodule hubs**.

```
hubs_gcn <- get_hubs_gcn(maize_exp, gcn)
head(hubs_gcn)
##              Gene Module  kWithin
## 1 Zm00001d049674  black 39.01150
## 2 Zm00001d017047  black 38.35905
## 3 Zm00001d042308  black 35.27548
## 4 Zm00001d013252  black 35.17820
## 5 Zm00001d035201  black 34.89403
## 6 Zm00001d011992  black 34.30860
```

Finally, we can visualize the network topology with hubs highlighted.

```
edges_filtered <- get_edge_list(gcn, module="black", filter=TRUE)
## The correlation threshold that best fits the scale-free topology is 0.8
```

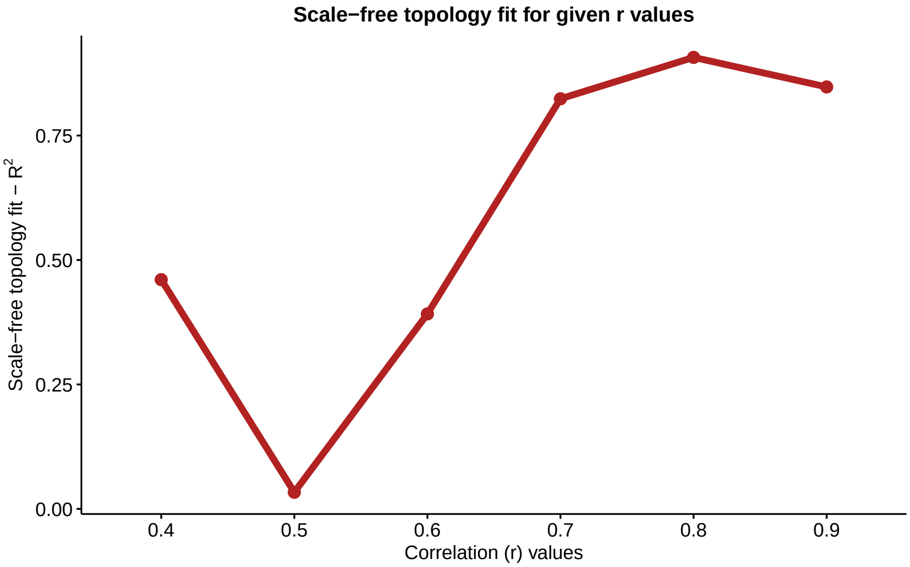

```
plot_gcn(edgelist_gcn = edges_filtered,
         net = gcn,
         color_by = "module",
         hubs = hubs_gcn)
```

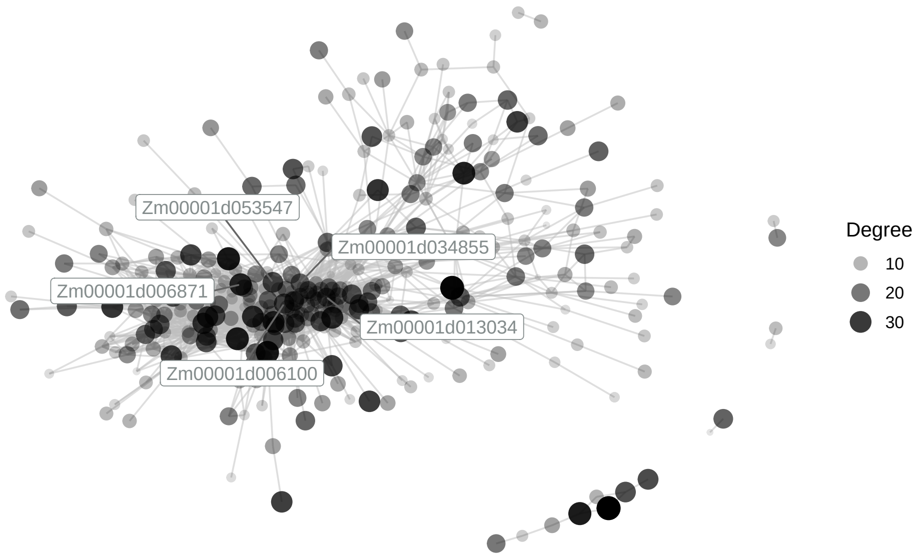

The highlighted genes are the top 5 hubs, which are likely the main genes controlling the biological processes associated with *black*.

### 5 GRN inference and analysis

Now, we will infer gene regulatory networks (GRNs), which are networks that depict interactions between regulators (e.g., transcription factors) and their target genes. Here, we will use the default method in `BioNERO`, which consists in inferring GRNs with 3 different algorithms (GENIE3, ARACNE, and CLR) and calculating average ranks for all edges across the algorithms. This principle, named *"wisdom of the crowds"*, ensures that the high-confidence edges we identify are not dependent on a single algorithm, but are a consensus across the three algorithms.

```
grn <- exp2grn(exp = maize_exp, regulators = zma_tfs$Gene)
## The top number of edges that best fits the scale-free topology is 178806
```

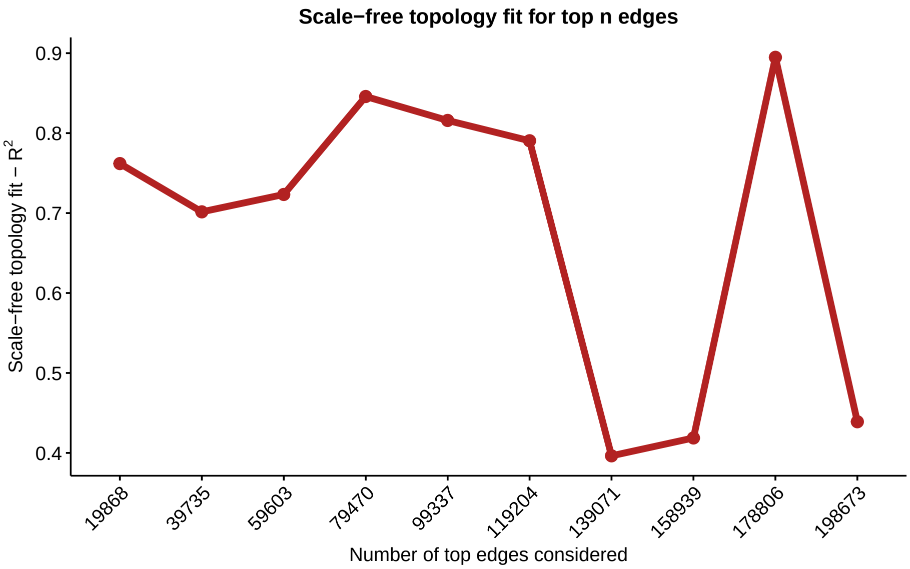

```
head(grn)
##             Regulator         Target
## 87212  Zm00001d023615 Zm00001d023616
## 115430 Zm00001d032032 Zm00001d019605
## 15798  Zm00001d005016 Zm00001d041962
## 89599  Zm00001d024547 Zm00001d024546
## 186655 Zm00001d049543 Zm00001d043023
## 15995  Zm00001d005016 Zm00001d053009
```

As we have the GRN as an edge list, we can find hubs (highly connected transcription factors) and visualize the network. For visualization, we will keep only the top 2000 edges ($R^2 > 0.8$) for a cleaner plot.

```
# Get hubs
grn_hubs <- get_hubs_grn(grn)
head(grn_hubs)
##             Gene Degree
## 1 Zm00001d034160   2174
## 2 Zm00001d021403   1765
## 3 Zm00001d042288   1630
## 4 Zm00001d007962   1503
## 5 Zm00001d003195   1450
## 6 Zm00001d017782   1448

# Visualize GRN
plot_grn(grn[1:2000, ])
```

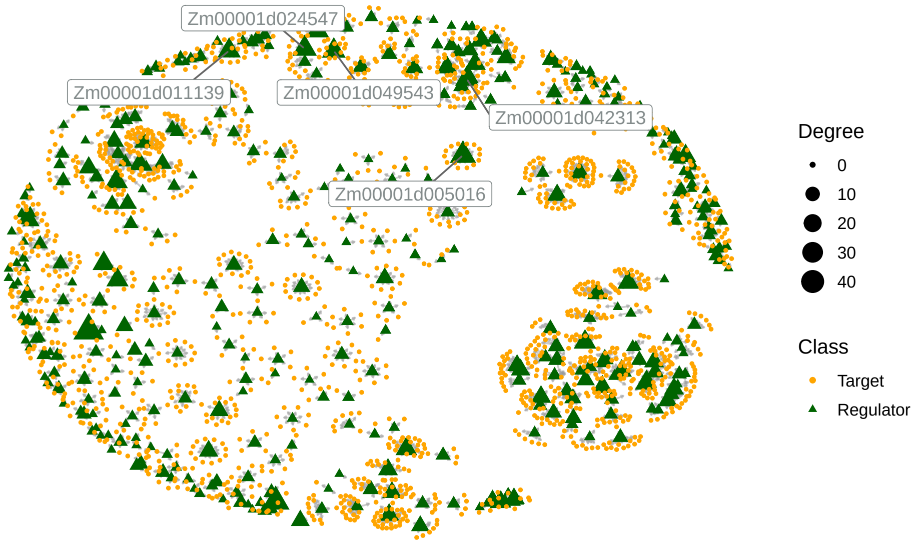

Additionally, we can use the *infomap* algorithm to detect communities in the GRN. Other algorithms are available in `detect_communities()`.

```
com <- detect_communities(grn)

# See number of communities
length(unique(com$mem))
## [1] 1

# See number of nodes per community
sort(table(com$mem), decreasing = TRUE)
##     1
## 14542
```

### 6 Module preservation

Module preservation is often used to study patterns of evolutionary conservation and divergence across transcriptomes. Here, we will compare rice and maize GCNs to explore how different these two monocot species are at the transcriptional level.

First of all, as gene IDs are different for the two species, we will need to collapse gene-level expression levels to orthogroup-level. Orthogroups were identified with OrthoFinder (Emms and Kelly 2015) and downloaded from the PLAZA 4.0 Monocots database (Van Bel et al. 2018).

```
head(orthogroups)
##                Family Species         Gene
## 1483 ORTHO04M000001     osa Os01g0100100
## 1484 ORTHO04M000001     osa Os01g0100200
## 1485 ORTHO04M000001     osa Os01g0100400
## 1486 ORTHO04M000001     osa Os01g0100500
## 1487 ORTHO04M000001     osa Os01g0100600
## 1488 ORTHO04M000001     osa Os01g0100700
```

```
# Store SummarizedExperiment objects in a list
zma_osa_list <- list(osa = rice_se, zma = maize_se)

# Collapse gene-level expression to orthogroup-level
ortho_exp <- exp_genes2orthogroups(zma_osa_list, orthogroups)

# Inspect new expression data
ortho_exp$osa[1:5, 1:5]
##                 SRX2273231 SRX2273206 SRX2273207 SRX2273209 SRX2273210
## ORTHO04M000001    2.34671   2.200730   1.893160    4.30026   3.908320
## ORTHO04M000002    3.19967   3.604760   1.108080    3.45536   2.921740
## ORTHO04M000003    4.93292   4.153930   2.681865    6.30378   5.533045
## ORTHO04M000004    4.00248   2.709920   2.216910    5.14441   4.141320
## ORTHO04M000005    5.22924   3.931355   3.183590    6.94365   6.575990
ortho_exp$zma[1:5, 1:5]
##                 SRX3141064 SRX3141035 SRX3141068 SRX3141040 SRX3141075
## ORTHO04M000001  3.6225900  3.4888800  3.5685100 0.39244800  0.4063130
## ORTHO04M000002  0.0395668  0.0384109  0.0000000 0.12220400  0.1335680
## ORTHO04M000003  0.1811055  0.2652720  0.0000000 0.34912950  0.3340505
## ORTHO04M000004  0.1481995  0.1841155  0.1227985 0.58693200  0.6624990
## ORTHO04M000005  0.0000000  0.0000000  0.0000000 0.08869925  0.0506705
sapply(ortho_exp, nrow)
##   osa   zma
## 12981 14813
```

Now, we will preprocess both expression sets with `exp_preprocess()`. We will only keep orthogroups with median expression levels >2.

```
ortho_exp <- lapply(ortho_exp, exp_preprocess, min_exp = 2)
## Number of removed samples: 12
## Number of removed samples: 5

# Check orthogroup number
sapply(ortho_exp, dim)
##       osa  zma
## [1,] 6990 7854
## [2,]  253  111
```

Now that row names are comparable, we will infer GCNs iteratively for each set.

```
# Calculate SFT power
power_ortho <- lapply(ortho_exp, SFT_fit,
                      cor_method="pearson",
                      net_type = "signed hybrid")
##    Power SFT.R.sq   slope truncated.R.sq mean.k. median.k. max.k.
## 1      3 0.000689  0.0157         -0.144 1190.00   1290.00 2330.0
## 2      4 0.137000 -0.2940          0.251  781.00    822.00 1760.0
## 3      5 0.279000 -0.5560          0.494  529.00    539.00 1360.0
## 4      6 0.386000 -0.7640          0.639  369.00    362.00 1060.0
## 5      7 0.449000 -0.9190          0.728  263.00    247.00  842.0
## 6      8 0.509000 -1.0500          0.796  191.00    170.00  675.0
## 7      9 0.552000 -1.1700          0.836  141.00    120.00  546.0
```

```
## 8     10 0.584000 -1.2700          0.866  105.00     84.60  445.0
## 9     11 0.606000 -1.3600          0.886   79.70     60.60  365.0
## 10    12 0.617000 -1.4300          0.901   61.10     43.70  302.0
## 11    13 0.635000 -1.5100          0.914   47.30     31.70  251.0
## 12    14 0.658000 -1.5500          0.929   36.90     23.30  209.0
## 13    15 0.678000 -1.5900          0.938   29.10     17.20  176.0
## 14    16 0.691000 -1.6300          0.947   23.10     12.80  148.0
## 15    17 0.709000 -1.6500          0.955   18.50      9.62  125.0
## 16    18 0.724000 -1.6700          0.960   14.90      7.25  106.0
## 17    19 0.736000 -1.6800          0.964   12.00      5.51   90.5
## 18    20 0.747000 -1.6800          0.965    9.81      4.18   77.4
## No power reached R-squared cut-off, now choosing max R-squared based power
##    Power SFT.R.sq  slope truncated.R.sq mean.k. median.k. max.k.
## 1      3    0.456 -0.875          0.651  486.00   419.000 1410.0
## 2      4    0.562 -1.150          0.743  278.00   208.000  987.0
## 3      5    0.631 -1.300          0.811  170.00   110.000  714.0
## 4      6    0.671 -1.410          0.849  109.00    62.100  530.0
## 5      7    0.700 -1.490          0.876   72.80    36.700  402.0
## 6      8    0.727 -1.530          0.901   50.00    22.700  310.0
## 7      9    0.742 -1.560          0.916   35.20    14.300  242.0
## 8     10    0.763 -1.580          0.932   25.30     9.310  191.0
## 9     11    0.781 -1.590          0.942   18.60     6.240  153.0
## 10    12    0.799 -1.580          0.951   13.90     4.220  123.0
## 11    13    0.806 -1.600          0.956   10.50     2.930  101.0
## 12    14    0.815 -1.600          0.958    8.02     2.020   83.9
## 13    15    0.816 -1.630          0.960    6.22     1.430   70.2
## 14    16    0.825 -1.620          0.962    4.87     1.030   59.1
## 15    17    0.840 -1.610          0.966    3.85     0.749   50.0
## 16    18    0.852 -1.590          0.968    3.08     0.554   42.5
## 17    19    0.887 -1.550          0.982    2.48     0.408   36.4
## 18    20    0.919 -1.520          0.991    2.01     0.307   31.2

# Infer GCNs
gcns <- lapply(seq_along(power_ortho), function(n) {
  return(exp2gcn(ortho_exp[[n]],
           SFTpower = power_ortho[[n]]$power,
           cor_method = "pearson",
           net_type = "signed hybrid",
           module_merging_threshold = 0.9))
  })
## ..connectivity..
## ..matrix multiplication (system BLAS)..
## ..normalization..
## ..done.
## ..connectivity..
## ..matrix multiplication (system BLAS)..
## ..normalization..
## ..done.

length(gcns)
```

```
## [1] 2
```

Finally, we can calculate module preservation statistics with the NetRep algorithm (Ritchie et al. 2016), which is based on non-parametric permutation analyses.

```
# Using rice as reference and maize as test
pres <- module_preservation(ortho_exp,
                            ref_net = gcns[[1]],
                            test_net = gcns[[2]],
                            algorithm = "netrep")
## [2021-04-10 14:57:23 -03] Validating user input...
## [2021-04-10 14:57:23 -03]   Checking matrices for problems...
## [2021-04-10 14:57:25 -03] Input ok!
## [2021-04-10 14:57:25 -03] Calculating preservation of network subsets from
##                           dataset "osa" in dataset "zma".
## [2021-04-10 14:57:25 -03]   Pre-computing network properties in dataset
##                             "osa"...
## [2021-04-10 14:57:27 -03]   Calculating observed test statistics...
## [2021-04-10 14:57:27 -03]   Generating null distributions from 1000
##                             permutations using 1 thread...
##
##
    0% completed.
    1% completed.
    1% completed.
    2% completed.
    2% completed.
    3% completed.
    4% completed.
    4% completed.
    5% completed.
    5% completed.
    6% completed.
    7% completed.
    7% completed.
    8% completed.
    8% completed.
    9% completed.
   10% completed.
   10% completed.
   11% completed.
   11% completed.
   12% completed.
   13% completed.
   13% completed.
   14% completed.
   14% completed.
   15% completed.
   16% completed.
   16% completed.
   17% completed.
```

```
          17% completed.
          18% completed.
          19% completed.
          19% completed.
          20% completed.
          20% completed.
          21% completed.
          22% completed.
          22% completed.
          23% completed.
          23% completed.
          24% completed.
          25% completed.
          25% completed.
          26% completed.
          26% completed.
          27% completed.
          28% completed.
          28% completed.
          29% completed.
          29% completed.
          30% completed.
          31% completed.
          31% completed.
          32% completed.
          32% completed.
          33% completed.
          34% completed.
          34% completed.
          35% completed.
          35% completed.
          36% completed.
          37% completed.
          37% completed.
          38% completed.
          38% completed.
          39% completed.
          40% completed.
          40% completed.
          41% completed.
          41% completed.
          42% completed.
          43% completed.
          43% completed.
          44% completed.
          44% completed.
          45% completed.
          46% completed.
          46% completed.
          47% completed.
          47% completed.
```

```
            48% completed.
            49% completed.
            49% completed.
            50% completed.
            50% completed.
            51% completed.
            52% completed.
            52% completed.
            53% completed.
            53% completed.
            54% completed.
            55% completed.
            55% completed.
            56% completed.
            56% completed.
            57% completed.
            58% completed.
            58% completed.
            59% completed.
            59% completed.
            60% completed.
            61% completed.
            61% completed.
            62% completed.
            62% completed.
            63% completed.
            64% completed.
            64% completed.
            65% completed.
            65% completed.
            66% completed.
            67% completed.
            67% completed.
            68% completed.
            68% completed.
            69% completed.
            70% completed.
            70% completed.
            71% completed.
            71% completed.
            72% completed.
            73% completed.
            73% completed.
            74% completed.
            74% completed.
            75% completed.
            76% completed.
            76% completed.
            77% completed.
            77% completed.
            78% completed.
```

```
   79% completed.
   79% completed.
   80% completed.
   80% completed.
   81% completed.
   82% completed.
   82% completed.
   83% completed.
   83% completed.
   84% completed.
   85% completed.
   85% completed.
   86% completed.
   86% completed.
   87% completed.
   88% completed.
   88% completed.
   89% completed.
   89% completed.
   90% completed.
   91% completed.
   91% completed.
   92% completed.
   92% completed.
   93% completed.
   94% completed.
   94% completed.
   95% completed.
   95% completed.
   96% completed.
   97% completed.
   97% completed.
   98% completed.
   98% completed.
   99% completed.
  100% completed.
  100% completed.
##
## [2021-04-10 15:00:14 -03]   Calculating P-values...
## [2021-04-10 15:00:15 -03]   Collating results...
## [2021-04-10 15:00:16 -03] Done!
## None of the modules in osa were preserved in zma.
```

As we have seen, none of the modules in rice were preserved in maize. The non-preservation of coexpression modules is likely due to different experimental conditions and biological variation between the two sets. For instance, there are stress-related samples in maize that do not have corresponding samples in rice. This is the main caveat in cross-species network comparison, as the difference between networks can be due to different experimental designs, not to intrinsic divergence between species. Thus, choosing which samples to include in each expression set is arguably the most important step for network comparison analyses, and it should be considered carefully.

### Session info

```
## R Under development (unstable) (2021-03-07 r80079)
## Platform: x86_64-pc-linux-gnu (64-bit)
## Running under: Ubuntu 16.04.7 LTS
##
## Matrix products: default
## BLAS:   /usr/local/lib/R/lib/libRblas.so
## LAPACK: /usr/local/lib/R/lib/libRlapack.so
##
## locale:
##  [1] LC_CTYPE=pt_BR.UTF-8       LC_NUMERIC=C
##  [3] LC_TIME=pt_BR.UTF-8        LC_COLLATE=en_US.UTF-8
##  [5] LC_MONETARY=pt_BR.UTF-8    LC_MESSAGES=en_US.UTF-8
##  [7] LC_PAPER=pt_BR.UTF-8       LC_NAME=C
##  [9] LC_ADDRESS=C               LC_TELEPHONE=C
## [11] LC_MEASUREMENT=pt_BR.UTF-8 LC_IDENTIFICATION=C
##
## attached base packages:
## [1] stats     graphics  grDevices utils     datasets  methods   base
##
## other attached packages:
##  [1] forcats_0.5.1    stringr_1.4.0    dplyr_1.0.5      purrr_0.3.4
##  [5] readr_1.4.0      tidyr_1.1.3      tibble_3.1.0     ggplot2_3.3.3
##  [9] tidyverse_1.3.0  BioNERO_0.99.8   BiocStyle_2.19.1
##
## loaded via a namespace (and not attached):
##   [1] readxl_1.3.1                backports_1.2.1
##   [3] circlize_0.4.12             Hmisc_4.5-0
##   [5] plyr_1.8.6                  igraph_1.2.6
##   [7] splines_4.1.0               GENIE3_1.13.3
##   [9] BiocParallel_1.25.5         GenomeInfoDb_1.27.10
##  [11] ggnetwork_0.5.8             sva_3.39.0
##  [13] digest_0.6.27               foreach_1.5.1
##  [15] htmltools_0.5.1.1           GO.db_3.12.1
##  [17] fansi_0.4.2                 magrittr_2.0.1
##  [19] checkmate_2.0.0             memoise_2.0.0
##  [21] cluster_2.1.1               doParallel_1.0.16
##  [23] limma_3.47.12               openxlsx_4.2.3
##  [25] ComplexHeatmap_2.7.9        fastcluster_1.1.25
##  [27] Biostrings_2.59.2           annotate_1.69.2
##  [29] modelr_0.1.8                matrixStats_0.58.0
##  [31] jpeg_0.1-8.1                colorspace_2.0-0
##  [33] ggrepel_0.9.1               rvest_1.0.0
##  [35] blob_1.2.1                  haven_2.3.1
##  [37] xfun_0.22                   jsonlite_1.7.2
##  [39] crayon_1.4.1                RCurl_1.98-1.3
##  [41] genefilter_1.73.1           impute_1.65.0
##  [43] survival_3.2-10             iterators_1.0.13
##  [45] glue_1.4.2                  gtable_0.3.0
##  [47] zlibbioc_1.37.0             XVector_0.31.1
```

```
##  [49] GetoptLong_1.0.5            DelayedArray_0.17.10
##  [51] car_3.0-10                  shape_1.4.5
##  [53] BiocGenerics_0.37.1         abind_1.4-5
##  [55] scales_1.1.1                edgeR_3.33.3
##  [57] DBI_1.1.1                   rstatix_0.7.0
##  [59] Rcpp_1.0.6                  xtable_1.8-4
##  [61] htmlTable_2.1.0             clue_0.3-58
##  [63] foreign_0.8-81              bit_4.0.4
##  [65] preprocessCore_1.53.2       Formula_1.2-4
##  [67] stats4_4.1.0                htmlwidgets_1.5.3
##  [69] httr_1.4.2                  RColorBrewer_1.1-2
##  [71] ellipsis_0.3.1              farver_2.1.0
##  [73] pkgconfig_2.0.3             XML_3.99-0.6
##  [75] dbplyr_2.1.0                nnet_7.3-15
##  [77] locfit_1.5-9.4              utf8_1.2.1
##  [79] dynamicTreeCut_1.63-1       labeling_0.4.2
##  [81] reshape2_1.4.4              tidyselect_1.1.0
##  [83] rlang_0.4.10                AnnotationDbi_1.53.1
##  [85] cellranger_1.1.0            munsell_0.5.0
##  [87] tools_4.1.0                 cachem_1.0.4
##  [89] cli_2.4.0                   generics_0.1.0
##  [91] RSQLite_2.2.5               broom_0.7.6
##  [93] evaluate_0.14               fastmap_1.1.0
##  [95] yaml_2.2.1                  fs_1.5.0
##  [97] RhpcBLASctl_0.20-137        knitr_1.31
##  [99] bit64_4.0.5                 zip_2.1.1
## [101] KEGGREST_1.31.1             nlme_3.1-152
## [103] xml2_1.3.2                  compiler_4.1.0
## [105] rstudioapi_0.13             curl_4.3
## [107] png_0.1-7                   ggsignif_0.6.1
## [109] minet_3.49.0                reprex_1.0.0
## [111] statmod_1.4.35              geneplotter_1.69.0
## [113] stringi_1.5.3               ps_1.6.0
## [115] lattice_0.20-41             Matrix_1.3-2
## [117] vctrs_0.3.7                 networkD3_0.4
## [119] pillar_1.5.1                lifecycle_1.0.0
## [121] BiocManager_1.30.10         GlobalOptions_0.1.2
## [123] cowplot_1.1.1               data.table_1.14.0
## [125] bitops_1.0-6                GenomicRanges_1.43.4
## [127] R6_2.5.0                    latticeExtra_0.6-29
## [129] bookdown_0.21               network_1.16.1
## [131] gridExtra_2.3               rio_0.5.26
## [133] IRanges_2.25.6              codetools_0.2-18
## [135] assertthat_0.2.1            SummarizedExperiment_1.21.2
## [137] DESeq2_1.31.18              rjson_0.2.20
## [139] withr_2.4.1                 S4Vectors_0.29.13
## [141] GenomeInfoDbData_1.2.4      intergraph_2.0-2
## [143] mgcv_1.8-34                 hms_1.0.0
## [145] parallel_4.1.0              grid_4.1.0
## [147] rpart_4.1-15                NetRep_1.2.4
## [149] rmarkdown_2.7.4             carData_3.0-4
```

```
## [151] MatrixGenerics_1.3.1       Cairo_1.5-12.2
## [153] ggpubr_0.4.0               ggnewscale_0.4.5
## [155] lubridate_1.7.10           Biobase_2.51.0
## [157] WGCNA_1.70-3               base64enc_0.1-3
```

### References

10 Emms, David M., and Steven Kelly. 2015. "OrthoFinder: solving fundamental biases in whole genome comparisons dramatically improves orthogroup inference accuracy." *Genome Biology* 16 (1): 1–14. https://doi.org/10.1186/s13059-015-0721-2.

Jin, J, F Tian, D C Yang, Y Q Meng, L Kong, J Luo, and G Gao. 2017. "PlantTFDB 4.0: toward a central hub for transcription factors and regulatory interactions in plants." *Nucleic Acids Res* 45 (D1): D1040–45. https://doi.org/10.1093/nar/gkw982.

Ritchie, Scott C., Stephen Watts, Liam G. Fearnley, Kathryn E. Holt, Gad Abraham, and Michael Inouye. 2016. "A Scalable Permutation Approach Reveals Replication and Preservation Patterns of Network Modules in Large Datasets." *Cell Systems* 3 (1): 71–82. https://doi.org/10.1016/j.cels.2016.06.012.

Shin, Junha, Harald Marx, Alicia Richards, Dries Vaneechoutte, Dhileepkumar Jayaraman, Junko Maeda, Sanhita Chakraborty, et al. 2020. "A network-based comparative framework to study conservation and divergence of proteomes in plant phylogenies." *Nucleic Acids Research*, 1–23. https://doi.org/10.1093/nar/gkaa1041.

Van Bel, Michiel, Tim Diels, Emmelien Vancaester, Lukasz Kreft, Alexander Botzki, Yves Van De Peer, Frederik Coppens, and Klaas Vandepoele. 2018. "PLAZA 4.0: An integrative resource for functional, evolutionary and comparative plant genomics." *Nucleic Acids Research* 46 (D1): D1190–96. https://doi.org/10.1093/nar/gkx1002.
